# Sulfidogenic Bacteria and the Risk of Colorectal Cancer

**DOI:** 10.64898/2026.07.14.738438

**Authors:** Darren Lee, Nicholas Ollberding, Qing Duan, Jordan Kharofa

**Affiliations:** Department of Radiation Oncology, University of Cincinnati, Cincinnati, OH; Division of Biostatistics and Epidemiology, Cincinnati Children’s Hospital Medical Center and Department of Pediatrics, University of Cincinnati College of Medicine, Cincinnati, OH

**Keywords:** Sulfidogenic bacteria, sulfur-metabolizing enzymes, colorectal cancer, gut microbiome

## Abstract

**Background:** Sulfur-rich dietary patterns have been associated with colorectal cancer risk. Certain gut bacteria metabolize dietary sulfur compounds into hydrogen sulfide, which can damage the intestinal epithelium and promote carcinogenesis. We evaluated whether sulfur-metabolizing enzymes are enriched in colorectal cancer and whether associations differ by age.

**Results:** Across eleven metagenomic cohorts, several sulfur-metabolizing enzymes were more prevalent in individuals with colorectal cancer than in healthy controls. Enzymes involved in sulfur reduction and detoxification, including sulfolactaldehyde reductase, peptide-methionine sulfoxide reductase, dimethylsulfoxide reductase, glutathione transferase, hydroxyacylglutathione hydrolase, and arylsulfatase, were consistently enriched in colorectal cancer. In contrast, enzymes involved in cysteine and methionine synthesis were more common in healthy controls. Age-stratified analyses showed minimal effect modification. *Fusobacterium nucleatum*, *Intestinimonas butyriciproducens*, and *Bilophila wadsworthia* carried more sulfur-metabolizing genes in colorectal cancer samples. Early-onset colorectal cancer samples were enriched for these genes in *Citrobacter*, *Klebsiella*, and *Raoultella*, whereas healthy controls showed greater representation of detoxification-associated genes in *Bifidobacterium* species.

**Conclusions:** Individuals with colorectal cancer harbor a greater abundance of sulfur-metabolizing enzymes, supporting a potential role for microbial conversion of dietary sulfur into hydrogen sulfide in colorectal carcinogenesis. Because these genes are distributed across many taxa, microbial function may better explain disease risk than individual species. Associations were consistent across age groups, suggesting that prolonged sulfur-rich dietary exposure may foster a microbial environment capable of generating carcinogenic metabolites. Microbial sulfur metabolism may therefore represent a modifiable pathway for prevention and further mechanistic study.

## Background

Epidemiologic studies have revealed higher rates of colorectal cancer (CRC) in individuals who adhere to enriched sulfur microbial diets. Through sulfur metabolism, certain bacterial species may produce genotoxic hydrogen sulfide (H_2_S) and these metabolic byproducts may contribute to diet mediated carcinogenesis(1–3). Although the incidence and death rate from CRC has declined in patients older than 50 years of age with the adoption of screening, CRC incidence in young patients has been increasing in the United States (4,5) and world-wide (6–11) with the highest increase noted in rectal cancer. Studies evaluating the fecal metagenome have revealed enrichment in distinct species in CRC patients when compared to healthy controls ^12–1512–15^,provoking the hypothesis that microbial features may, in part, explain the relative increase in young colon and rectal cancer incidences and could also underpin diet associations with colorectal cancer at large. However, potential microbial-host interactions responsible are not well described. We have previously demonstrated that sulfur metabolizing species, including *Bilophilia wadsworthia* and *Alistipes putredinis*, are differentially enriched in CRC patient samples age 20-49 relative to healthy controls (16). However, it has become apparent that sulfur metabolizing genes may be enriched across various taxa comprising the human microbiome (17).

Wolf et al. recently showed that sulfur metabolizing enzymatic pathways are present in more commensal bacterial species than previously realized(17). The metabolic function of the microbiome may reflect the expression of conserved genes across various species. Sulfur metabolizing species may produce H_2_S via multiple pathways including metabolism of inorganic sulfate directly or through metabolism of organic sulfur amino acids like cysteine, methionine, and taurine(17). Dissimilatory sulfite reductase is the enzyme involved in the final step of H_2_S production by sulfate or taurine respiration. In a cohort of 514 metagenomic samples from the Human Microbiome Project, Wolf et al. detected dissimilatory sulfite reductases in 18% of samples distributed across various phyla(17). Thus, there is strong rationale to evaluate the association of CRC and specific sulfur metabolizing genes and enzyme-catalyzed reactions agnostic to a given species genome for all patients, as well as to assess if these genes and reactions are enriched in young onset CRC. Nguyen et al. have previously identified notable bacterial genes involved in sulfur metabolism in a prior work(1) and have derived the EC numbers for relevant sulfur metabolizing enzymes of interest.

We hypothesized that sulfidogenic bacteria act as mediators of CRC carcinogenesis and potentially contribute to the increase in early onset rectal through dietary metabolism of sulfur to H_2_S. In addition, critical insights may be obtained by identifying the relative abundance of sulfur metabolizing genes in the genomes of specific bacterial species across the age spectrum in patients with or without CRC. In this study, we evaluate associations of sulfur metabolizing genes agnostic to bacterial species with CRC overall and as a function of age.

## Method

### Data acquisition

Analyses was performed using metagenomic data obtained via the curated MetagenomicData package (18)in the R software environment as published in our prior work (19). In our prior published metanalysis, all available studies were reviewed to identify samples containing colorectal cancer patients and healthy controls with eleven datasets identified meeting these criteria(1,12–15,20–24). This resulted in 692 patients with colorectal cancer and 609 healthy controls. The curatedMetagenomicData package provides uniformly processed human microbiome data from previously published and publicly available data. The methods for generating the required data have been described previously in detail (25–27). For our analysis, we obtained the relative abundance of all available UniRef90 gene families using the function returnSamples("gene_families", count = F) then assign Enzyme Commission (EC) numbers to gene families using the utility mapping accessory files provided in the HUMAnN3 software program. We focused on the 52 sulfur metabolizing EC reactions described by Nguyen et al (1)..

### Statistical analysis

Sample metadata and EC number relative abundances were converted to phyloseq objects for analysis using the phyloseq package in R. Means with standard deviations and frequencies with percents were used to describe participant demographic and clinical characteristics according to cancer status. For our primary analyses, the relative abundance for each enzyme-catalyzed reaction were converted to presence/absence due to the expected high proportion of zeros and low abundances as performed in our previously published work.

A two-stage individual patient data meta-analysis (IPDMA) approach were used to obtain summary odds ratios (OR) and 95% confidence intervals (CI) for colorectal cancer case status according to the presence of each enzyme catalyzed reaction. Logistic regressions were used to obtain log odds and standard errors (SE) for each study separately. Model terms included an indicator for enzyme presence/absence, age, gender, and body mass index (BMI). Age and BMI were modeled using restricted cubic spline terms with three knots placed at the 10^th^, 50^th^, and 90^th^ percentiles as recommend by Harrell(28). Summary ORs were obtained using the metafor package in R and estimated via restricted maximum likelihood (REML) with the Hartung-Knapp estimate of variance. Two-stage IPDMA with adjustment for gender, CRC status, and body mass index were used to obtain conditional summary ORs for a 10-year increase in age at the time of sample collection according to each enzyme presence overall and by cancer status.

Interaction ORs assessing the difference in the effect of each enzyme on CRC case status according to age < 50 years at onset (i.e., young onset) or older were obtained by pooling the log odds and SE for the cross-product terms for age (binary) and the enzyme in the second stage meta-analysis. Secondary analyses were conducted modeling the relative abundance as a linear term in the two-stage IPDMA framework described above. Generalized linear mixed effects regression (GLMER) were also used to model and visualize the prevalence of enzymes as a non-linear function of age. Estimates were obtained using GLMER with logit link function to model the binomially distributed responses; a study-specific random intercept; and fixed-effect terms for age, gender, BMI, cancer status, and the age by cancer status interaction. Age were modeled using restricted cubic spline terms with the number and placement of knots based on the model AIC. Models were fitted using the glmmTMB package in R and conditional effect plots generated using the sjPlot package to visualize sulfur metabolizing enzyme prevalence as function of age and cancer status.

The stratified UniRef90 gene families output from HUMAnN3 as provided by the CuratedMetagenomic package provide the relative contribution/abundance of each species included in the ChocoPhlAn reference database for each gene. We utilized this stratified dataset to first map gene families to enzyme-catalyzed reactions and then to quantify the relative abundance, as well as the presence/absence, of each species to each of the 52 enzymes of interest. We performed this for all samples and then according to cancer status (e.g., case and control) and age. Descriptive statistics including numbers and proportions with Wilson 95% CIs and stacked bar plots were used to describe and visualize the contribution of each species to the abundance or detection of each enzyme.

## Results

### Differential Expression of Sulfur-Metabolizing Enzymes in Colorectal Cancer

To understand the association of sulfur metabolism and CRC, we compared the sulfur-metabolizing enzyme gene relative abundance in CRC patients and healthy controls (Figure 1A). Several sulfur-metabolizing enzymes demonstrated significant differential abundance in CRC compared to controls. In particular, Sulfolactalehyde 3-reductase, oxidoreductases acting on a sulfur group of donors with NAD+ or NADP+ as acceptor, peptide-methionine (S)-S-oxide reductase, respiratory dimethylsulfoxide reductase, glutathione transferase, hydroxyacylglutathione hydrolase, and arylsulfatase were markedly upregulated in CRC. In contrast, cysteine synthase, O-acetylhomoserine aminocarboxypropyltransferase, and carbon-sulfur lyases were more abundant in controls compared to CRC patients.

**Figure 1.**
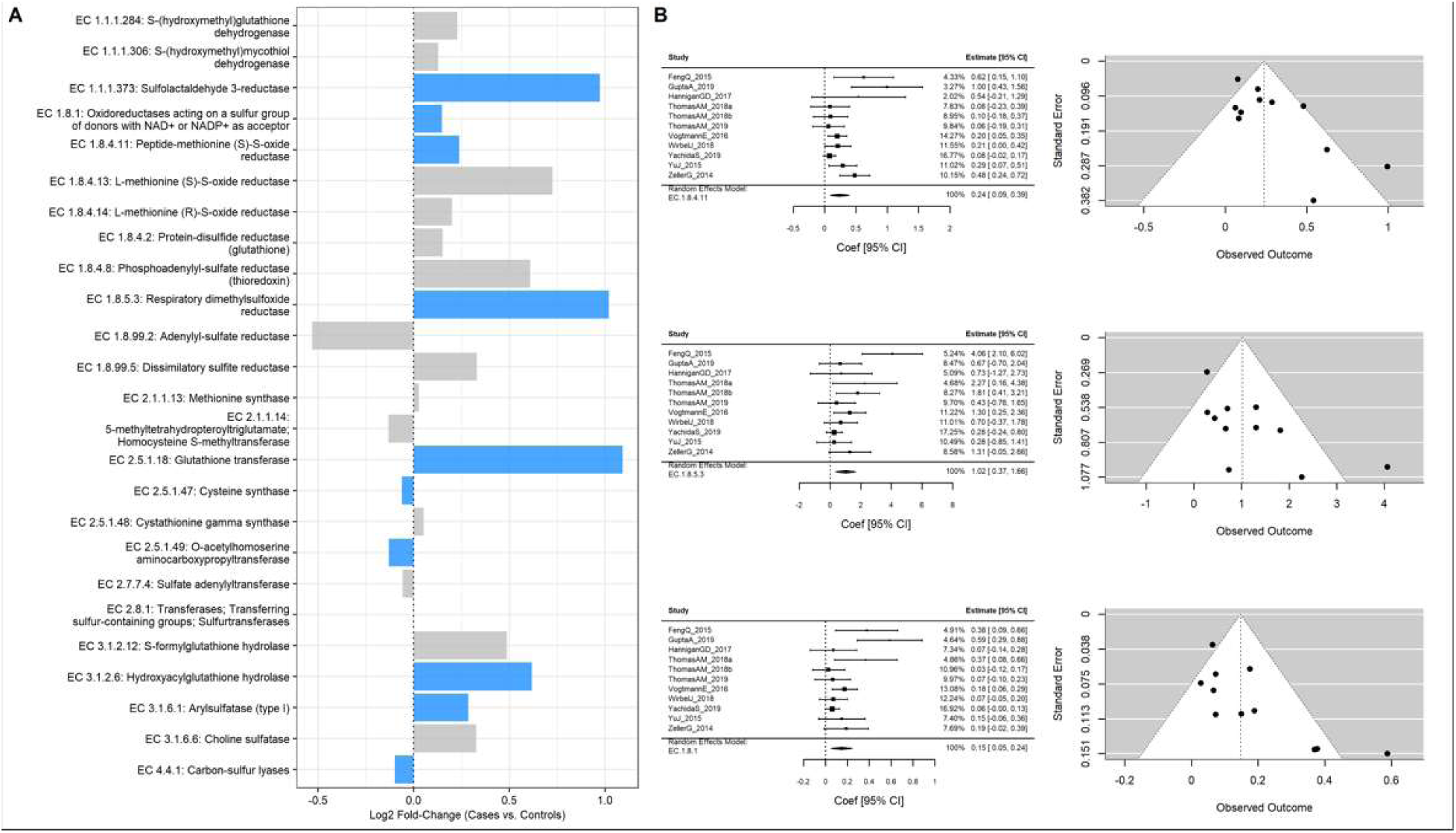
Differential abundance of sulfur-metabolizing enzymes in all CRC and controls. Colored bars in the differential abundance plot highlight gene families with p-values < 0.05. Forest plots estimate the pooled effect size of each study at 95% confidence interval. Funnel plots provide a visual representation of small-study effects and detect the presecne of publication bias.

To further quantify the strength and consistency of these associations, forest plots were generated for the top differentially expressed enzymes (Figure 1B). Across independent datasets, peptide-methionine (S)-S-oxide reductase, Sulfolactalehyde 3-reductase, respiratory dimethylsulfoxide reductase, and oxidoreductases acting on a sulfur group of donors with NAD+ or NADP+ as acceptor exhibited reproducible directionality and statistically significant effect estimates. Funnel plots showed no evidence of publication or sampling bias.

We next assessed whether age modified the association between sulfur-metabolizing enzymes and colorectal cancer (CRC) status (Figure 2A). The differential abundance between cases and control samples for individuals under 50 when compared to individuals 50 and above, adjusting for age, gender, and BMI (interaction with age<50) Age-stratified analyses revealed distinct metabolic signatures: O-acetylhomoserine aminocarboxypropyltransferase abundance increased with age, whereas S-formylglutathione hydrolase was preferentially reduced in younger patients. However, there were little material differences in the relative abundance of sulfur metabolizing enzymes as a function of age. Forest plots for the top age-modulated enzymes (Figure 2B) revealed reproducible directionality and effect sizes for the top differentially expressed enzymes, while corresponding funnel plots showed no evidence of major asymmetry or publication bias.

**Figure 2.**
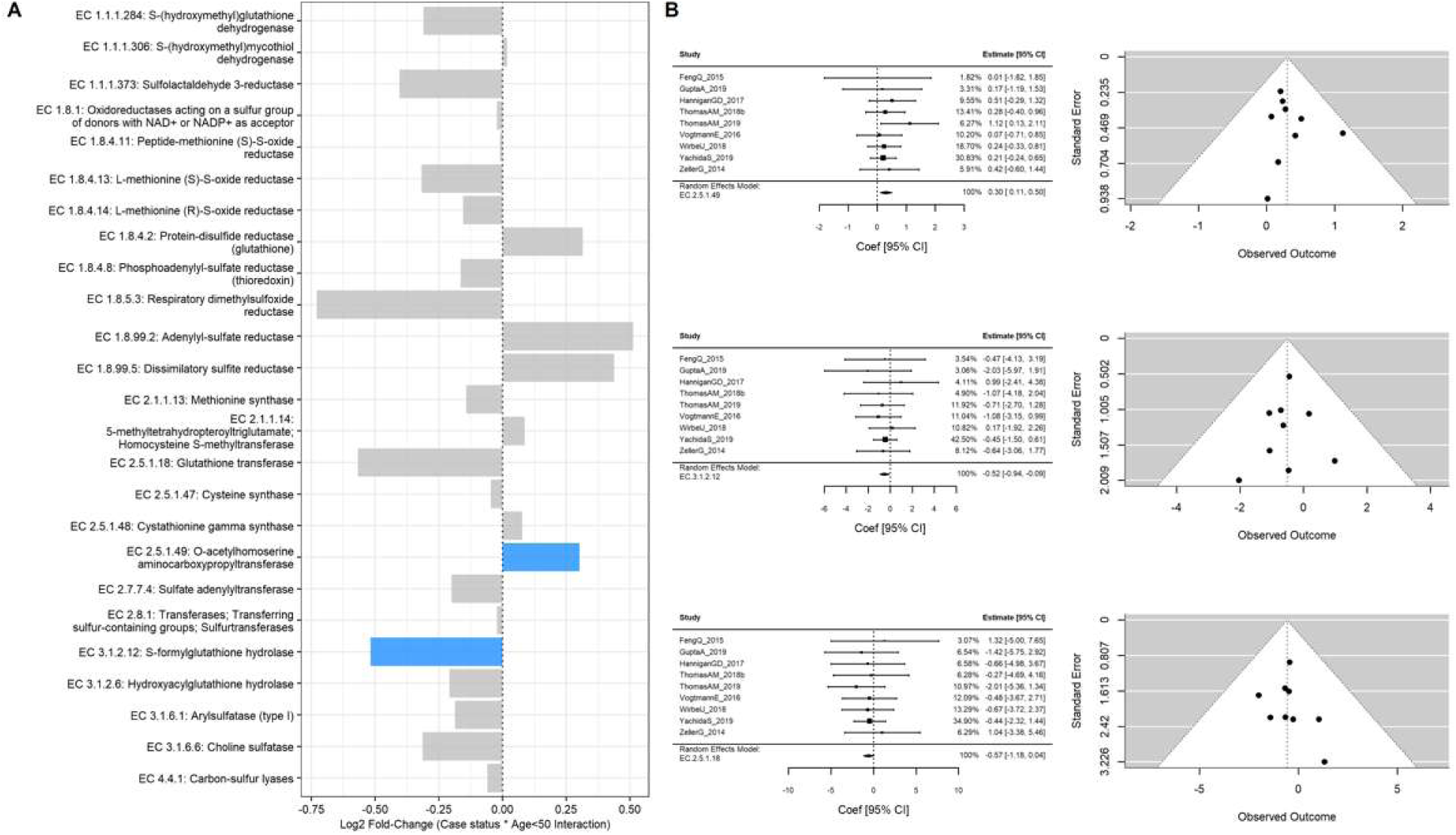
Differential abundance of sulfur-metabolizing enzymes in young-onset and older-onset CRC. Colored bars in the differential abundance plot highlight gene families with p-values < 0.05. Forest plots estimate the pooled effect size of each study at 95% confidence interval. Funnel plots provide a visual representation of small-study effects and detect the presecne of publication bias.

### Species-Specific Patterns of Sulfur-Metabolizing Genes

We examined the relative abundance of sulfur-producing genes in different species implicated in CRC, we compared the prevalence of these genes among All CRC patients, those with early-onset CRC (< 50 years), late-onset CRC (≥ 50 years), and healthy controls (Figure 3). The species of interest were selected based on prior known association with CRC and sulfur metabolism[29]. Species assessed included *Fusobacterium nucleatum, Firmicutes bacterium, Bilophila wadsworthia, Intestinimonas butyriciproducens, Holdemania filiformis, and Alistipes putredinis*. The prevalence of the sulfur metabolizing enzyme was calculated as the proportion of individuals within each group harboring a given gene. Across several species of interest, we observed the trend of increased prevalence of sulfur-metabolizing enzymes in the species collected from CRC patients compared to the same species when collected from healthy controls.

**Figure 3.**
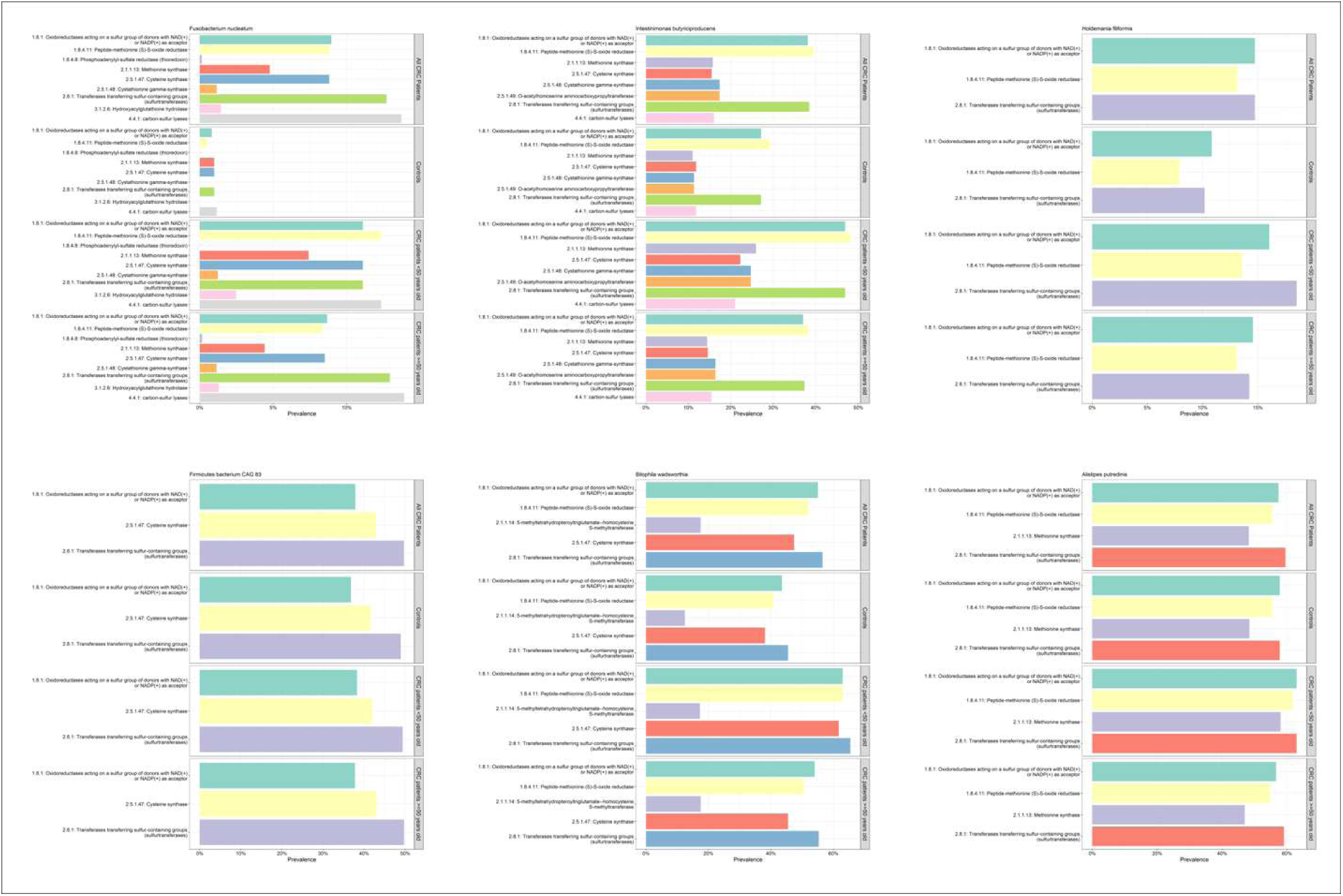
Relative abundance of sulfur-metabolizing genes of top sulfidenogenic bacterial species in healthy controls, young-onset CRC, older-onset CRC, and all CRC patients.

Multiple sulfur-metabolizing enzymes, including oxidorectases, methionine- and cysteine-synthesizing enzymes, sulfurtransferases, and carbon-sulfur lyases, were enriched in *Fusobacterium nucleatum and Intestimimonas butyriciproducens* derived from young-onset and older-onset CRC patients compared with healthy controls. *Intestimimonas butyriciproducens* exhibited a more pronounced enrichment in early-onset CRC.

Together, these patterns suggest that not only is the relative abundance of sulfur metabolizing genes greater in CRC compared to controls, but also that within the same species there may be increased prevalence of sulfur metabolizing enzyme presence when isolated from CRC vs controls. This suggests the possibility of strain level variability in sulfur metabolizing enzymes

### Taxonomic Distribution of Sulfur-Metabolizing Genes in CRC

We next evaluated the relative abundance of top 10 bacterial species harboring each sulfur-metabolizing genes significantly enriched in CRC patients compared to controls. For each gene, proportional abundances were normalized to the total abundance of all taxa carrying that gene. In younger-onset CRC, *Citrobacter freundii*, *Citrobacter youngae*, and *Klebsiella aerogenes* expressing sulfolactaldehyde 3-reductase were more abundant compared with both older-onset CRC and healthy controls. Similarly, younger-onset CRC samples expressing glutathione transferase showed enrichment in *Klebsiella aerogenes* and *Raoultella ornithinolytica* (Figure 4). In contrast, oxidoreductase-expressing communities in younger-onset CRC exhibited lower *Escherichia coli* abundance relative to older-onset CRC and healthy controls. Both younger- and older-onset CRC samples expressing hydroxyacylglutathione hydrolase were enriched in *Eisenbergiella tayi* compared with healthy controls. Conversely, S-(hydroxymethyl)glutathione dehydrogenase expression was associated with higher abundance in healthy controls.

**Figure 4.**
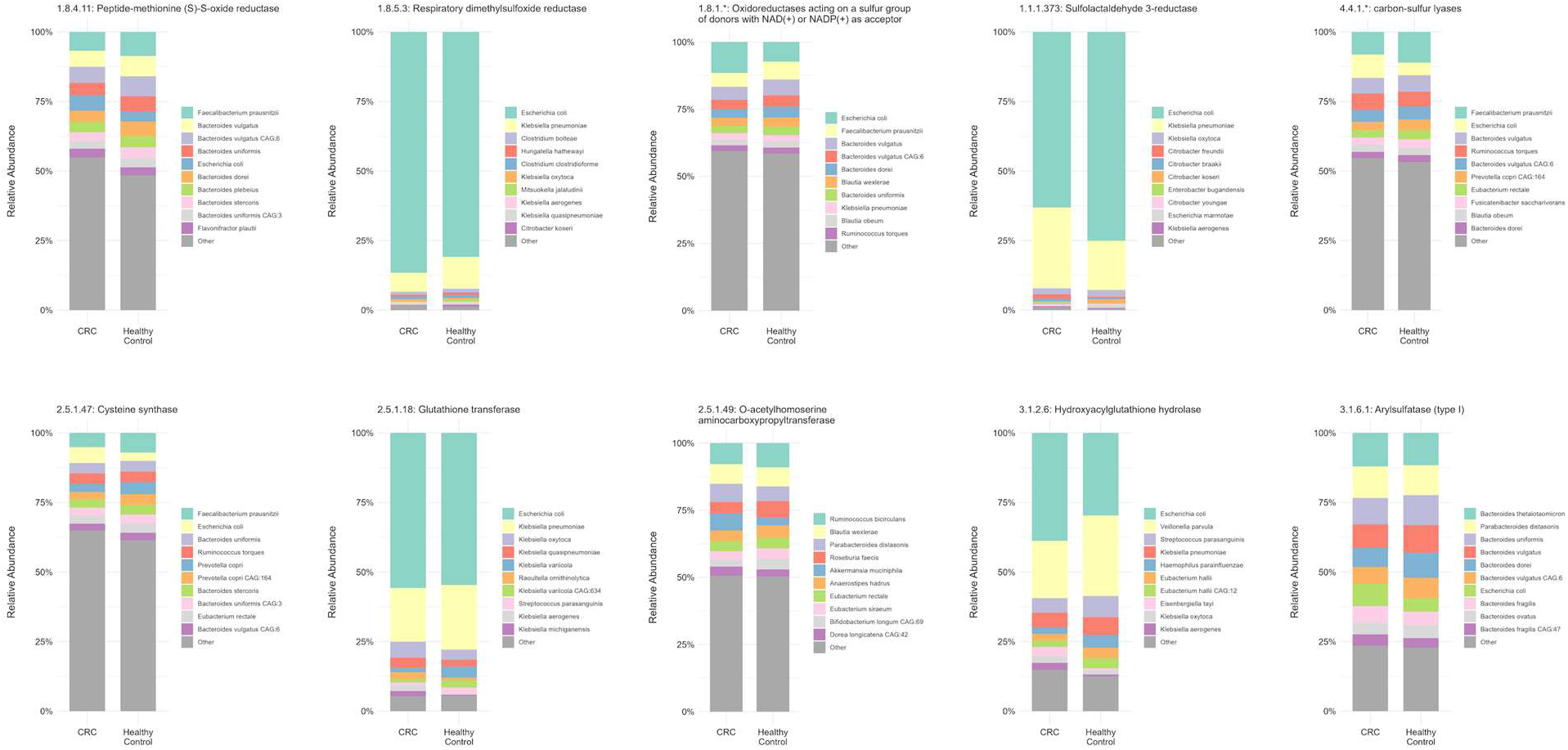
Relative abundance of sulfur-metabolizing enzymes in bacterial species enriched in CRC patients relative to controls.

Healthy controls displayed greater relative abundance of *Bifidobacterium bifidum* and *Bifidobacterium pseudocatenulatum* carrying S-(hydroxymethyl)glutathione dehydrogenase (Figure 4). CRC samples expressing S-(hydroxymethyl)mycothiol dehydrogenase were enriched in *Actinomyces odontolyticus*, *Enterococcus casseliflavus*, *Rothia aeria*, and *Corynebacterium simulans*, but showed reduced *Rothia dentocariosa* abundance. CRC samples expressing sulfoacetaldehyde reductase were enriched in *Klebsiella michiganensis* and *Klebsiella quasipneumoniae*, while those expressing dimethyl sulfide:cytochrome C₂ reductase were enriched in *Klebsiella oxytoca*; by contrast, healthy controls exhibited enrichment of *Citrobacter amalonaticus*. CRC expressing dissimilatory sulfite reductase were enriched in *Desulfovibrio fairfieldensis* and *Desulfovibrio legallii*. Finally, protein-S-isoprenylcysteine O-methyltransferase expression was associated with enrichment of *Candida albicans* and *Giardia intestinalis* in CRC samples.

## Discussion

The primary finding from this study demonstrates a higher relative abundance of sulfur-metabolizing enzymes in patients with colorectal cancer compared to healthy controls. This observation supports growing evidence that the metabolic activity of the gut microbiome, rather than its microbial composition, may be a key link between diet and colorectal carcinogenesis. It is plausible that sulfur-metabolizing enzymes interact with dietary sulfur compounds (e.g., taurine, cysteine, and sulfate preservatives) to produce carcinogenic metabolic byproducts such as hydrogen sulfide. H_2_S has been observed to have carcinogenic potential by directly damaging DNA in mammalian cells or indirectly by disrupting the normal colonic mucosal layer (29–32).The data presented herein justify further investigation into the role of microbial sulfur metabolism and its modulation by diet as part of the diet–microbiome–host axis in CRC development.

Large prospective cohort studies and systematic reviews have shown that long-term adherence to a Western diet increases the risk of CRC(33–35). However, the exact underlying mechanism for these associations was not previously established. Rather than direct carcinogensis, it is possible that host microbial features may act as intermediaries producing carcinogenic metabolites from dietary substrates. A Western diet that high in animal protein, fat, and processed foods, and low in fiber is associated with high sulfur microbial diet. Adherence to a sulfur microbial diet (rich in processed meats, soft drinks, and liquor but low in vegetables and legumes) has been associated with an increased risk of CRC in multiple epidemiologic cohorts (1–3). In a prospective cohort of 307 healthy men from the Men’s Lifestyle Evaluation Study, investigators prospectively measured dietary patterns to evaluate correlation of fecal metagenomics with adherence to a sulfur microbial diet(2). *Bilophilia wadsworthia* was one of two notable species enriched in patients with a sulfur microbial diet. In a separate study of 30,818 women less than 50 years old, investigators examined dietary patterns and association with early onset CRC. They noted that higher sulfur microbial diet scores in early adulthood were associated with increased risk of early-onset conventional adenomas (1). Stool correlates to examine microbial features were unavailable.

Recently, CRC screening guidelines have changed from 50 years to 45 years of age (36). However, the incidence in the United States and other western populations in patients younger than age 45 years old continues to increase (4,37). Using the Surveillance, Epidemiology, and End Results (SEER) database, investigators noted a 1-3.2% annual increase in colorectal cancer in patients aged 20-39 years old since the mid-1980s (4). This increase in Young Onset CRC is not uniform across the United States. Notably, the incidence is higher in the UCCC catchment and neighboring states relative to other parts of the United States.

In a previous, we examined the relative abundance of all species differentially enriched in young onset CRC compared to controls and older patients with CRC (16). Notably, we observed enrichment of *Bilophilia wadsworthia* and *Alistipes putredinis* in young onset CRC relative to both older CRC patients and healthy controls. This finding is intriguing given their role in sulfur metabolism and production of H_2_S and had not been previously demonstrated in young onset CRC. We did not observe a significant interaction between age and sulfur-metabolizing enzyme abundance. One explanation may be that if diet–microbiome interactions primarily drive sulfur metabolism, these relationships exist consistently across the age spectrum. In other words, the rising incidence of young-onset CRC may partly reflect earlier or prolonged exposure to sulfur-rich diets that favor sulfur-metabolizing microbiomes capable of producing carcinogenic metabolites.

Supporting this, epidemiologic data from large women’s health cohorts demonstrate that higher adherence to sulfur microbial dietary patterns is associated with increased risk of early-onset CRC and adenoma formation(1). Collectively, these findings strengthen the rationale for further mechanistic studies examining sulfur metabolism as a modifiable microbial pathway linking diet, microbiome composition, and colorectal cancer development across age groups.

In this multi-cohort metagenomic analysis, we found that sulfur-metabolizing enzymes are consistently more abundant in individuals with colorectal cancer than in healthy controls, supporting the hypothesis that microbial sulfur metabolism may contribute to colorectal carcinogenesis. The enrichment of enzymes involved in hydrogen sulfide production and sulfur compound reduction across diverse bacterial taxa suggests that microbial species makeup, rather than the presence of any single pathogenic species, may be the more relevant driver of disease biology. Although we observed minimal interaction between age and enzyme prevalence, species-level analyses revealed distinct microbial patterns in younger-onset colorectal cancer, including greater representation of Klebsiella, Citrobacter, and Raoultella species carrying sulfur-metabolizing genes. These findings raise the possibility that early or prolonged exposure to sulfur-rich dietary patterns may shape microbial metabolic activity long before cancer develops.

Taken together, our results highlight microbial sulfur metabolism as a potentially modifiable pathway linking diet, microbiome composition, and colorectal cancer risk. Future mechanistic studies are warranted to define how sulfur-metabolizing enzymes influence mucosal integrity, genomic stability, and host inflammatory responses, as well as to evaluate whether diet-based or microbiome-targeted interventions can reduce the burden of colorectal cancer across age groups.

## Declarations

### Data Availability Statement

All metagenomic data analyzed in this study were obtained from the curatedMetagenomicData package, a publicly available, curated resource accessible through Bioconductor’s ExperimentHub. No original sequence data were generated in this study. The curatedMetagenomicData package is available at https://waldronlab.io/curatedMetagenomicData/ and via Bioconductor (https://bioconductor.org/packages/curatedMetagenomicData). Data access and use are described in: Pasolli E, et al. Accessible, curated metagenomic data through ExperimentHub. *Nat Methods.* 2017;14(11):1023–1024. https://doi.org/10.1038/nmeth.4468.

### Ethics Statement

This study was conducted using publicly available, de-identified metagenomic data obtained from the curatedMetagenomicData repository. No human subjects were enrolled, and no identifiable private information was accessed. In accordance with 45 CFR §46.104(d), this study qualifies for exemption from Institutional Review Board (IRB) oversight

### Authors’ contributions

N.O., Q.D., and J.K. conducted the data processing and statistical analyses. Manuscript draft was prepared by D.L. and J.K. All authors approved the final version.

## Acknowledgements

The authors thank the contributing investigators whose publicly available metagenomic datasets made this analysis possible, as well as the University of Cincinnati Department of Radiation Oncology and Cincinnati Children’s Hospital Division of Biostatistics and Epidemiology for supporting the computational resources used in this study.

